# Synthetic Cellular Immunity based on CAR repertoire for Human Immunotherapy

**DOI:** 10.1101/868653

**Authors:** Wenyan Fu, Chuqi Wang, Zetong Ma, Tian Li, Fangxing Lin, Ruixue Mao, Changhai Lei, Jian Zhao, Shi Hu

## Abstract

The immune surveillance hypothesis is regarded as the intellectual underpinning of cancer immunology. We find that engineered T cell library displaying CAR constructs, where the antibody repertoires took the place of the antigen-recognition region, has the capacity to recognize nonself-antigens and showed antigen dependent clonal expanding and persistence. This process coupled with long-lasting anti-tumour effect in a broad spectrum of epithelial tumours. Moreover, CAR-T library showed robust immunologic memory and maintenance of diversity to recognize mutated or evolved tumours in both immunodeficiency and immunoresponsive murine tumor models. Large CAR library was further developed in logic-gated NK-92 cells to develop an off-the shelf therapeutic library. Thus, the biological behaviour of synthetic cell library is highly accordance with the immune surveillance function, which proved the way to rebuilt functional MHC unrestricted cellular immunity in vitro using synthetic method.

## Introduction

Over the past decade, significant progress has been made in modern immunotherapy driven by targeted therapies that inhibit tumour angiogenesis and intrinsic drivers of cancer cell growth. These immunomodulatory therapies enhance host antitumour immunity and the basis of genetically engineered autologous or allogeneic immune cells expressing chimeric antigen receptors (CARs) or T-cell receptors (TCRs).

However, all current immunotherapies rely on the development of “specific targets”. In one respect, identification of the underlying cell-autonomous, gene-driven targets of tumorigenesis led to the development of clinically important blockers, such as EGFR or HER2 blockers, that resulted in profound, but often not durable, tumour responses in genetically defined patient populations. Another approach, exporting the targets involved with protective tumour immunity, has become a hot research area, most notably the “immune checkpoint” antibodies that reverse the action of negative regulators of T cell function that has led to durable clinical responses in subsets of patients with various tumour types. Adoptive CAR-based therapies, although showing very encouraging efficiency, also depend on the construction of a specific target binding domain, which is usually achieved by development of antigen-recognition regions in the form of a single-chain variable fragment (scFv) or a binding receptor/ligand in the extracellular domain. In fact, the theory of “targeting a specific target” creates a dilemma in the context of cellular plasticity and evolution, not to mention genetic or epigenetic heterogeneity of the disease, at every level from the molecule to the human population.

The human immune system never functions by depending on “specific targets”. Human immune cells act as natural defenders to protect the body from invading pathogens and to provide surveillance and respond to tumorigenesis. One scientific theory that describes the functions and processes adopted by lymphocytes to handle a wide variety of antigens that attack the body^1^ is Burnet’s clonal selection theory. Introduced and promoted by the Frank Macfarlane Burnet, the clonal selection theory explains the formation process of diverse antibodies and immunoglobulin-like proteins in the initial stages of the immune response. In the immune system, some lymphocytes with diverse immunoglobulin-like proteins are tolerant of self-tissues, while others are intolerant of self-tissues. However, notably, only cells that tolerate self-tissue survive the embryonic stage of development. In addition, the introduction of a nonself tissue leads to the development of lymphocytes that include nonself tissues as part of their self-tissues^2,3^. Frank Macfarlane Burnet proposed the clonal selection theory to explain and examine the functions of lymphocytes in the immune system and to assess how they respond to specific antigens that invade the body. Moreover, the theory provides a basis for understanding how the immune system responds to infections and how B and T lymphocytes are usually selected to destroy particular antigens.

The immunosurveillance hypothesis posits that because of the ability of host immune cells to recognize and destroy nascent malignancies, tumours arise at more frequent intervals than they are detected^4^. This hypothesis is further evidenced by the link of immune incompetence with an increased incidence of cancers^5^. Tumour-associated antigens (TAAs), which are mutated self-proteins expressed by some tumours, are capable of eliciting a tumour-specific immune response ^6^; however, the immune unresponsiveness hypothesis suggests that failure of the immune system to recognize TAAs leads to the evolution of manifested cancerous disease^7^. Accumulated ^8^evidence has led to the development of the cancer immunoediting hypothesis, which unifies the tumour protective immunosurveillance and tumour-promoting immune unresponsiveness aspects of immunity^9^. A corollary of this hypothesis, that the immune system shapes tumour immunogenicity, indicates that, to be effective, cancer immunotherapies must involve large numbers of high-quality effectors that can eliminate tumours and tumour-induced immune suppressors.

T cells, known to be key effectors of antitumour effects, are attractive candidates for cancer immunotherapy^10^. Theoretically, reconstruction of a healthy, tumour-responsive immune system in patients results in recognition and destruction of the TAA-expressing tumour cells. There is also suggestive evidence for this theory. Transplantation of haematopoietic cells from allogeneic donors supplemented with mature donor T cells can cure minimal residual disease and prevent leukaemia relapse ^11^. In a path-breaking trial by Hans Kolb and colleagues ^12^, adoptively transferred peripheral blood mononuclear cells (PBMCs) derived from the peripheral blood of donors were used to treat three patients who suffered from chronic myeloid leukaemia (CML) relapse after bone marrow transplantation. All three patients achieved haematologic and cytogenetic remission that persisted up to 7 months, thus documenting the first trial of allogeneic T-cell-based immunotherapy. Of note, two of the three patients treated with allogeneic PBMCs developed graft-versus-host disease (GVHD).

CAR-T cells, in contrast to conventional effector T cells, can be activated through the recognition of antigens irrespective of the MHC presentation. Interestingly, recent reports show that CAR-T cells originating from a single clone have the potential to show antitumour activity accompanied by complete remission in patients. We hypothesize that a CAR-T cell repertoire (or T cell CAR display) that has the potential to recognize a variety of nonself antigens can be directly used as a therapeutic agent. In this report, we designed and characterized an artificial bionic immune cell system based on synthetic receptor libraries. We postulate that, in a pre-existing group or population of genetically engineered lymphocytes, a particular antigen can exclusively activate its specific counter cell, enabling that specific cell to survive and multiply to destroy particular antigens. When the diversity of genetically engineered lymphocytes is sufficient to accurately and precisely recognize any pathogen or disease antigen, in theory, the cell population has the capacity to combat any disease in humans.

## Results

### A synthetic cell based CAR library

We first intend to construct a synthetic cell display library. In this library, the CAR library was first designed to displayed in T cells, where a ScFv library was fused to a CD8a hinge and transmembrane domain and the intracellular domains of human CD28 and CD3ζ (or z), consistent with second generation CAR design. A retroviral vector encoding inducible caspase-9 (iCASP9), IL15, and GFP separated by 2A sequences was further constructed (Fig. 1a). Engineered T cells with either an EGFR-specific or a HER2-specific CAR were characterized to evaluate the CAR design in our experiments. We confirmed the EGFR-CAR and HER2-CAR expression by staining with anti-Myc mAb (Fig. 1b). We then investigated the antitumour potential of the transduced T cells by standard ^51^Cr-release assays using MCF-7 cells (EGFR- and HER2-negative cells), MCF-7 EGFR cells (a derivative engineered to express EGFR), and the MCF-7 HER2 cells (a derivative engineered to express HER2). The MCF-7 cells and the derivative cells were characterized in our previous study^13^. CAR-T cells transduced with cetuximab scFv (termed CAR-T-CTX) efficiently lysed the EGFR-positive cells, that is, the MCF-7 EGFR cells, but did not kill the MCF-7 cells or the MCF-7 HER2 cells. On the other hand, the CAR-T cells transduced with trastuzumab scFv (termed CAR-T-TTZ) efficiently lysed the HER2-positive cells, that is, the MCF-7 HER2 cells, but not the MCF-7 cells or MCF-7 HER2 cells (Fig. 1c). To confirm activation-dependent IL15 production in T cells as reported previously, control T cells and different CAR-T cells were cultured the MCF-7 cells and the derivative cells. After 24 hours, IL15 concentration in culture media was determined by ELISA. Our data shows CAR-T cells produced significant amounts of IL15 only in the presence of antigen positive cells (fig. S1a).Moreover, postexposure to AP1903 caused significant apoptosis/necrosis of the engineered T cells with iCASP9 (fig. S1b).

**Figure 1.**
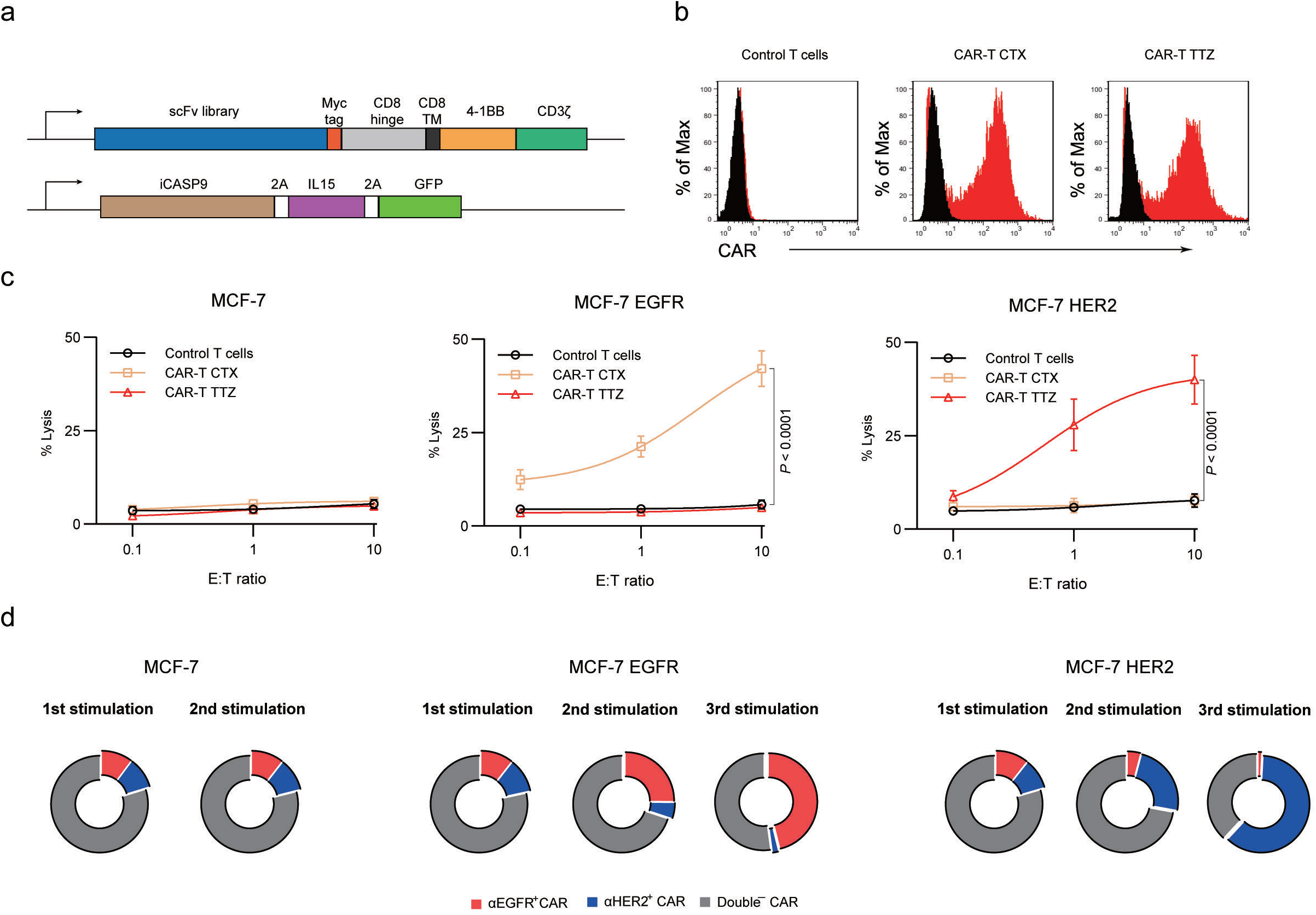
Design and characterization of a synthetic chimeric antigen receptor T cell library. **a.** Map of lentiviral constructs encoding the CAR library. **b.** Membrane-bound CAR expression. Forty-eight hours after retroviral transduction, the expression of CARs on human T cells was measured by staining with anti-MYC antibody, followed by flow cytometry analysis. T cells without transduction were used as negative controls. The histograms shown in black corresponds to the isotype controls, whereas the red histograms indicate positive fluorescence. **c.** Killing activity of the CAR-T cells in response to tumour cells. The cytotoxic activity of the CAR-T and control T cells against MCF-7 cells was assessed by a ^51^Cr-release assay at the indicated effector-to-target (E:T) ratios. **(d)** Growth dynamics of CAR-T library in the different co-culture methods cells were first stimulated with different MCF-7 cells for 24 or 72 hours. The addition of 10 nM AP1903 to the cultures induced apoptosis/necrosis of the CAR-T cells, as assessed by annexin-V staining in 4 independent experiments. The dimerizer did not induce apoptosis in the untransduced T cells.

For proof of concept, A CAR-T cell library of 10 control CAR constructs, which showed negligible binding to MCF-7 cells, termed as CAR-T_Control_ library (fig. S2). We confirmed the uniform expression of each of the 10 negative control CAR constructs on the cell surface of engineered T cells. Before conducting the experiments, a CAR-T cell display library was also developed in which CAR-T cells were mixed, and CAR-T CTX, CAR-T-TTZ and 8 other unique CAR-T cells were equally distributed in the population; this library was termed the CAR-T_CTX-TTZ_ library. To determine the dynamics of CAR composition, CAR-T_CTX-TTZ_ library were stimulated every 7 days with MCF-7 cells or MCF-7 derivatives. Prior to each stimulation the percentage of αEGFR^+^ CAR-T cells or αHER2^+^ CAR-T cells were analysed. Repeat stimulations with MCF7 EGFR cells or MCF-7 HER2 cells resulted in a notable enrichment of CAR-T CTX and CAR-T TTZ cells in comparison with other CAR-T cells (Fig. 1d), while there was no significant change of CAR composition of CAR-T_CTX-TTZ_ library co-cultured with MCF-7 cells after the first and second simulation. To test the synthetic cell based library in vivo, immunodeficient MCF-7, MCF-EGFR and MCF-HER2 xenograft NSG mice were maintained without treatment until a tumour of approximately 50 mm^3^ formed, and then, the mice were treated with CAR-T_CTX-TTZ_ library (1×10^6^ cells for each CAR construct containing T cells). For the MCF-7 xenograft mice, the tumours progressed rapidly, reaching a volume of 1,000 mm^3^ in fewer than 40 days. Interestingly, the mice that received the CAR-T_CTX-TTZ_ library showed significantly inhibited growth of the MCF-7 EGFR tumours or the MCF-7 HER2 tumours approximately two weeks after injection of CAR-T cells and the tumours showed complete regression at the end of the experiment, suggested the expansion of the target-specific CAR-T cells in these models (Fig. 2a). After the experiment, peripheral blood was collected from tumour-bearing mice and quantified for persistent human T cells as well as the frequencies of the different CAR-T cells. Notably, CAR^+^ T-cell counts in mice bearing MCF-7 EGFR or the MCF-7 HER2 tumours treated with CAR-T_CTX-TTZ_ library was significantly higher than in mice bearing MCF-7 tumours (Fig. 2b), indicating that antigen recognition drives the survival of the adoptively transferred T cells in *vivo*. Moreover, the enrichment of antigen-specific CAR-T cells were also observed in mice bearing MCF-7 EGFR or the MCF-7 HER2 tumours treated with CAR-T_CTX-TTZ_ library (Fig. 2c).

**Figure 2.**
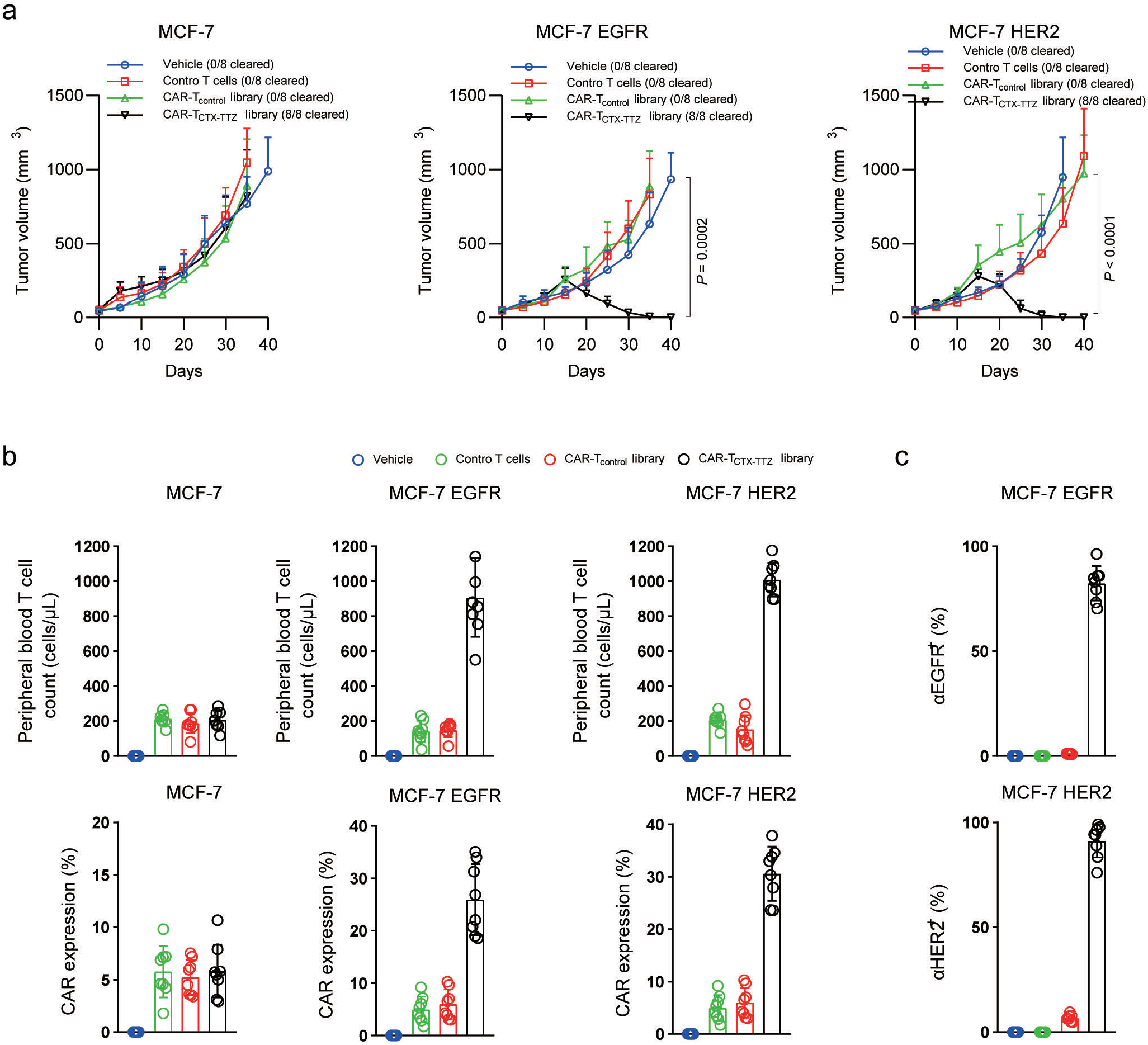
In vivo antitumour effect of the chimeric antigen receptor T cell library. **a.** Tumour volumes of different MCF-7 tumour xenografts after the indicated treatment. **b.** peripheral blood was collected and quantified for the absolute number of human T cells/ml of blood. **c.** antigen-specific CAR expression on human CAR T cells from peripheral blood of treated mice measured by flow cytometry.

To further explore our findings, we next aim to construct small-scale chimeric antigen receptor T cell library. Blood samples from 200 nonimmunized, healthy volunteers were collected to construct a nonimmunized human single-chain variable fragment (scFv) library. B lymphocyte cDNA encoding a variable fragment was used to construct a phage display scFv library that consisted of ~ 1×10^8^ individual colonies. The scFv gene corresponded to the size of the insert of more than 98% of the colonies. To confirm the heterogeneity of the individual clones from the library, we sequenced 50 randomly selected clones, and each clone showed a distinct scFv sequence. To prevent phage antibody library derived from human resource binding to mice tissues, we used phage that did not bind to systemic organs of mice as the source of the scFvs. Four rounds of in vivo phage screen were performed in the NSG mice to collect this “negative” phage population (Fig. 3a). The DNA sequences of the isolates were analysed, and the results confirmed that phage diversity was maintained, and this library termed libraryNSG. Previously, anti-EGFR CAR-T cells delayed the growth of SW480 colon tumours but unexpectedly induced the outgrowth of EGFR-negative tumour cells^14^; therefore, we used the SW480 cells in our study with the interest to find CARs has the potential to overcome resistance. A further four rounds of in vivo phage screen were performed in SW480 xenograft NSG mice injected with libraryNSG (Fig. 3b). Approximately 400 unique scFv sequences, indicative of tumour-binding scfvs, were identified in this manner and then use to construct a CAR-T cell library, termed as CAR-T_sw480_ library. As we used standard transfection methods to generate CAR-T cell libraries, the introduction of DNA encoding a repertoire of CAR genes has the potential to introduce multiple CAR genes into one cell. Next, the SW480 cells were injected subcutaneously into the NSG mice, and the mice were then injected (i.v.) with 1 × 10^7^ control T cells or CAR-T CTX cells when the tumour volume reached approximately 50 mm^3^ (Fig. 3c). Consistent with the findings of a previous report, tumour growth was significantly inhibited by the anti-EGFR-based cell therapy^14^. However, at approximately 3 weeks, the mice treated with the CAR-T CTX cells eventually succumbed to the tumour. The mice were then treated with 1× 10^6^ CAR-T_sw480_ libraries on day 25. Interestingly, approximately another three weeks after treatment, six of the eight mice receiving CAR-T_sw480_ library therapy showed significantly inhibited tumour growth, and the tumours showed complete regression two weeks later. After the experiments, all the remaining mice were sacrificed, and no αEGFR^+^ CAR-T cells were detected in any of the tested mice. The CAR-positive T cells were sorted and sequenced, and we were able to define five unique scFv clones, which were then used to construct to CARs, termed as CAR S1-S5. Combined treatment of CAR-T CTX cells and CAR-T S2 or CAR-T S3 cells in SW480 xenografts shows no sign of resistance (Fig. 4d). These two scFvs did not binds to EGFR therefore suggested binding to other tumour-antigens which have capacity to overcome resistance (fig. S3).

**Figure 3.**
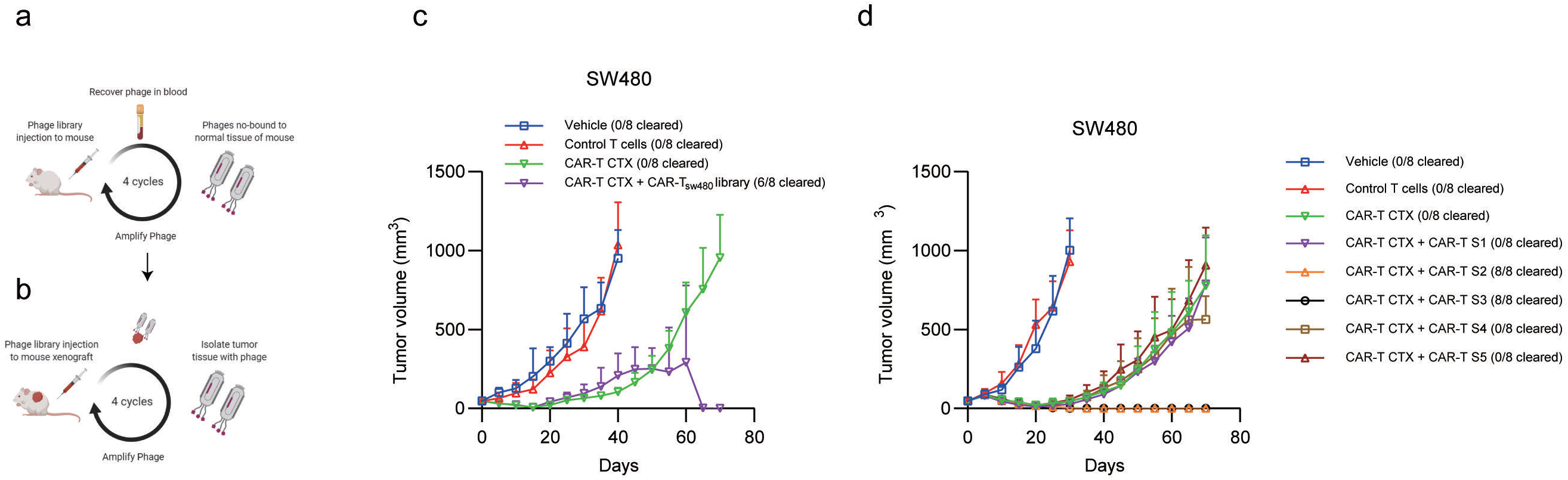
A Chimeric antigen receptor T cell library based on phage display technology. **a.** Negative selection: an in vivo phage display was performed to collect nonbinding phages to normal tissues in mice. **b.** After negative selection, specific scFvs homing to the cancer cells were identified by in vivo phage display in the NSG mice bearing SW480 xenograft cells. **c and d.** Tumour volumes of SW480 tumour xenografts after the indicated treatment.

**Figure 4.**
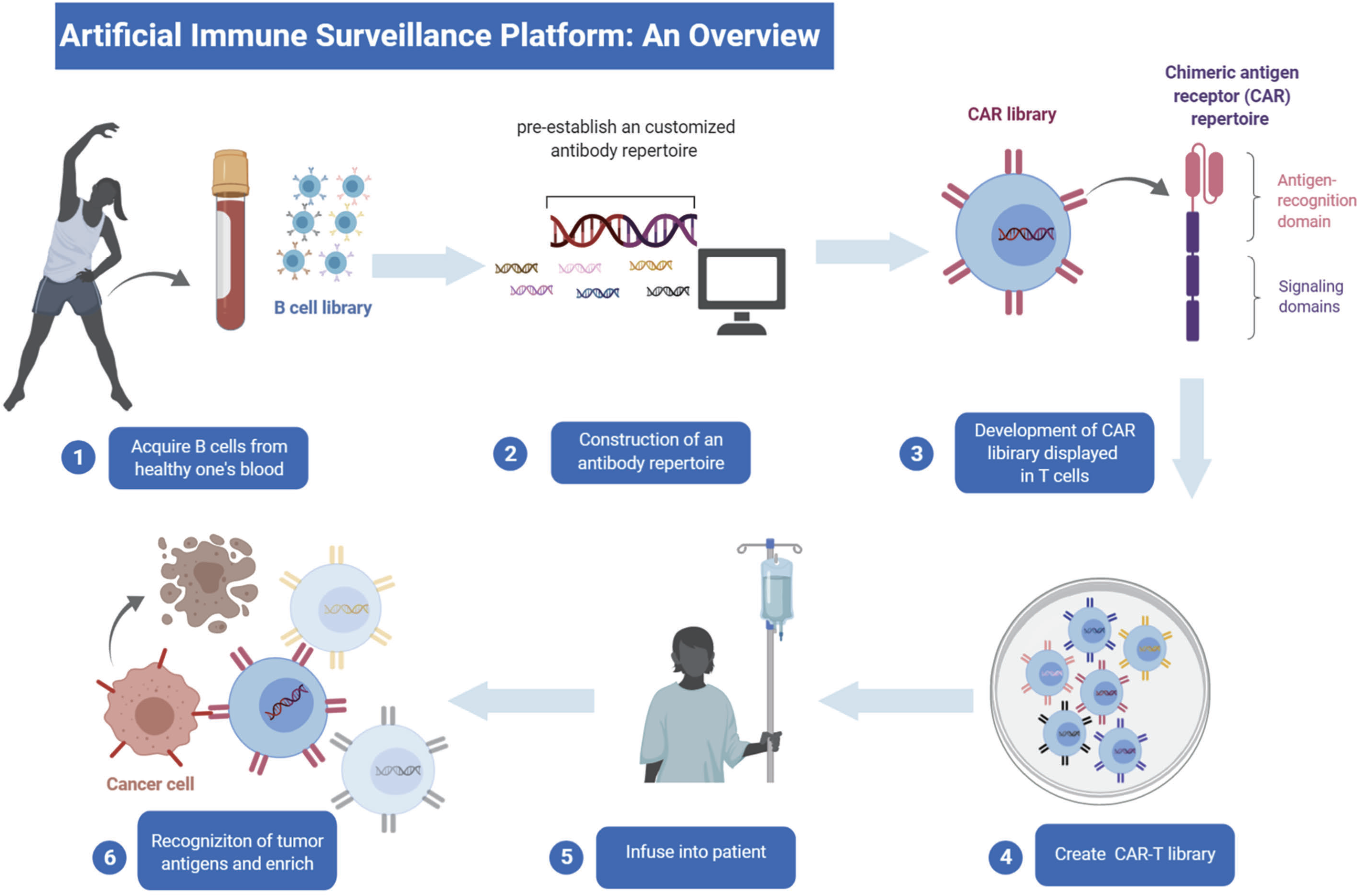
A Proposed Synthetic Immune Surveillance Platform.

### A Proposed Synthetic Immune Surveillance Platform

The observation that the chimeric antigen receptor T cell library have the capacity to recognize novel tumour antigens and this process coupled with the inhibition of tumour growth led us to form a bold idea: to develop an artificial immune surveillance platform using the CAR library (Fig 4). In this platform, a customized antibody repertoire is pre-established, which can be derived from a small number of antibodies binding to validated targets, natural naïve human antibody gene repertoires constructed from healthy donors’ B cells, or synthetic libraries & semi-synthetic libraries consisting of only synthetic sequences or a mixture of natural and synthetic antibody sequences, respectively. Because Naïve library, which is relatively close to the human antibody germ line and has low risk of immunogenicity, contain a very broad repertoire of antigen specificities theoretically covering every possible antigen, it is of interest and preferable in the platform for human therapeutic purpose. Naïve antibody libraries are constructed from rearranged V genes from B cells of non-immunized donors, i.e., the IgM repertoire. These antibodies have already undergone *in vivo* selection for functional B cell receptors, including a selection for low immunogenicity and low toxicity. Antibodies bind to allogeneic antigens can be further excluded before using such repertoires. The pre-established antibody repertoire was then used to construct synthetic receptor libraries displayed by engineered immune cells. The synthetic receptor can be activated and sustain the expansion and proliferation of engineered immune cells, moreover, mediate a specific biological activity, e.g. cellular killing effect for cancer therapeutic propose. Therefore, in this platform, the synthetic cell library have the captivity to recognize newly arising tumours through the expression of tumour specific neo-antigens on tumour cells and eliminate them, similarly to the homograft rejection and the “immunological surveillance mechanism”.

In this scenario, we first develop a CAR-T cell library containing 5×10^5^ individual constructs was developed based on the library_NSG_, containing candidates that showed no binding to systemic organs in NSG mice, as explained in the experiments described above, termed as CAR-T_NSG_ library. The antitumour efficacy of the CAR-T_NSG_ library in the NSG mice xenografted with SW480 cells were further investigated, notably, one of eight mice treated with CAR-T_NSG_ library tumours initially developed but completely regressed under therapy, with the mice being tumour-free latest by day 75, while we observed no signs of CAR-T cell expansion at the end of the experiments in the other seven mice in this treatment arm (Fig. 5a). One mice do response to the CAR-T_NSG_ library suggested that the diversity of the CAR library is enough to recognize tumour-antigen. Because each CAR clone was only displayed in a related small amount of T cells, we hypothesis the failure of enrichment of tumour specific CAR-T cells may be due to lack of long-term T-cell persistence.

**Figure 5.**
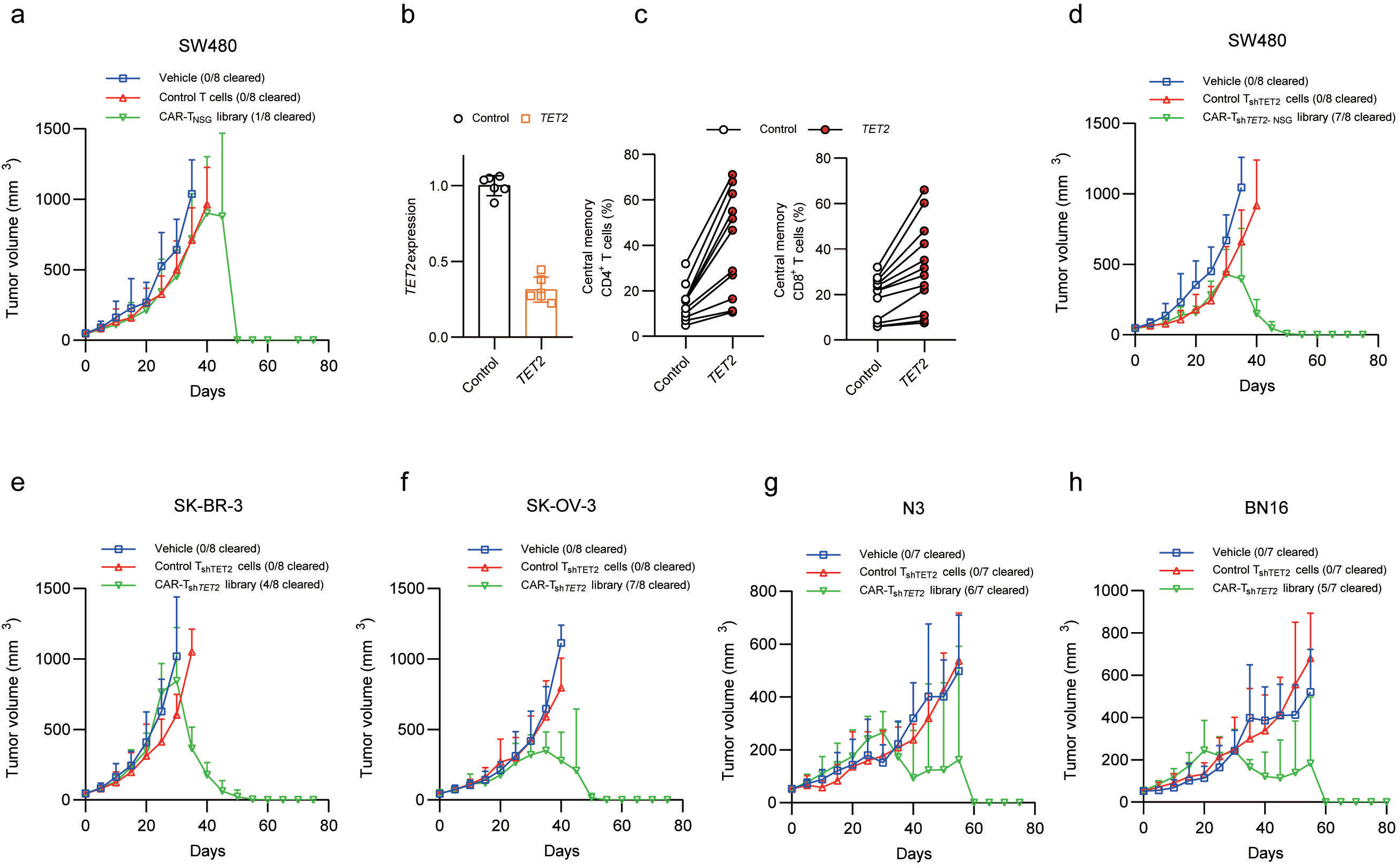
Characterization of a large scale chimeric antigen receptor T cell library. **a** Tumour volumes of SW480 tumour xenografts after the indicated treatment. **b.** *TET2* expression in the T cells transduced with a scrambled shRNA (control) or TET2 shRNA. **c.** Frequencies of central memory CD8^+^ T cells (left) and CD4^+^ T cells (right) following shRNA-mediated knockdown of *TET2* (n = 12; pooled results from 4 independent experiments). Tumour volumes of SW480 **(d)**, SK-BR-3 **(e)**, SK-OV-3 **(f)**, N3 **(g)**, BN16 **(h)** tumour xenografts after the indicated treatment.

A previous study showed that a single *TET2*-modified CAR-T cell had the capacity to induce leukaemia remission in human and that CAR-T cells exhibited a central memory phenotype^15^. We therefore used *TET2*-knockdown CAR-T cells (referred to as CAR-T_*shTET2*_ cells) to construct a synthetic cell library (Fig. 5b). The experimental knockdown of *TET2* in the T cells resulted in a central memory phenotype in the CD8^+^ and CD4^+^ T cells (Fig. 5c), and a CAR-T_sh*TET2-*NSG_ library containing 5×10^5^ individual constructs based on the library_NSG_ was also developed. The CAR-T_sh*TET2-*NSG_ library or parental T_sh*Tet2*_ cells were used to treat the mice bearing the established subcutaneous SW480 xenograft tumours. Our data showed that tumours grew rapidly in the control mice injected with medium or parental T_*shTET2*_ cells; the tumours treated with the CAR-T_sh*TET2-*NSG_ library did not show any signs of inhibition until approximately one month later, at which time, tumour growth was inhibited (Fig. 5d). Completely regressed tumours were observed in 7 of the 8 tumours in the mice receiving therapy, although one mice in the CAR-T_sh*TET2-*NSG_ library treatment group were sacrificed during the 7^th^ week because the tumours had become excessively large. Next, the potential for the CAR-T_sh*TET2-*NSG_ library to eradicate tumours in mouse xenografts was further examined. Mice xenografted with SK-BR-3 cancer cells serving as a models of HER2^+^ breast cancer were treated with a transfer of the CAR-T_sh*TET2-*NSG_ library or parental T_*shTET2*_ cells (Fig. 5e). Tumours in the mice treated with the CAR-T_sh*TET2-*NSG_ library were significantly inhibited and then eliminated compared with those in the mice treated with parental T_*shTET2*_ cells, although four mice in the CAR-T_sh*TET2-*NSG_ library treatment group were sacrificed during the 7^th^ and 8^th^ weeks for the outgrowth of tumours. The therapeutic effect of the CAR-T_sh*TET2-*NSG_ library was further examined in another model using SK-OV-3 cells, an ovarian HER2^+^ cancer cell line (Fig. 5f). Transfer of the CAR-T_sh*TET2-*NSG_ library significantly inhibited tumour growth approximately one month later, and the tumours were eradicated in 7 of 8 mice, with one of which were sacrificed for tumours that were excessively large.

We next utilized previously characterized patient-derived xenograft (PDX) models to further investigate the antitumor effects of CAR-T_sh*TET2-*NSG_ library. While the control T cell treated mice rapidly developed tumours, six of seven CAR-T_sh*TET2-*NSG_ library treated mice engraftment with the NSCLC tumour N3 displayed inhibition of tumour growth and finally free of tumour (Fig. 5g). The same experiments performed in a TNBC BN16 PDX model yielded similar results, completely regressed tumours were observed in five of the seven tumours in the mice receiving CAR-T_sh*TET2-*NSG_ library treatment (Fig. 5h). For the PDX models that experienced a complete response, we rechallenged these mice with a second implantation. The rechallenged mice received no additional therapy. Of note, the rechallenge N3 cells or BN16 cells were incapable of establishing a tumor mass in mice previously cured of either tumors, indicate that rechallenge tumors can be eliminated by the enrichment of tumor-specific CAR-T cells (fig. S4a and S4b). Next, we validated whether the CAR-T_sh*TET2*-NSG_ library maintain the targeting diversity after long-term tumor-free survival. We rechallenged the mice previously cured of N3 or BN16 tumor challenged by with the colorectal cancer cell line SW480 to test whether the CAR-T cell library retain its capacity to recognize a complete different tumour (fig. S4c and S4d). We observed that SW480 cells have the ability to engraft in the rechallenged mice, remarkably, we observed an inhibition of tumor growth about one month after rechallenge. Only one non-responder was observed in our study with significant tumor burden requiring euthanization within 50 days after tumor rechanllge. Taken together, these results indicate that CAR library displayed in T cells have the potential to enrichment tumor-specific CAR-T cells eliminating established tumor, and the diversity of the library can be maintained during long-term tumor-free survival.

### Development of a Naïve Mouse Synthetic Immune Surveillance Platform

A Naïve mouse scFv library constructed from the spleen cells of C57BL/6 mice with approximately 3 × 10^8^ unique clones were developed. The murine scFv library were further used to construct a CAR library in *Tet2*-knockdown murine T cells (T_sh*Tet2*_ cells) (fig. S5), termed as CAR-T_sh*Tet2*-C57_ cell library, and a cell library containing approximately 7× 10^6^ unique CAR clones was used for the further study selection. In the pre-study in our experiments, the engraftment of donor T cells was very low, we therefore employ pretreatment with cyclophosphamide (CPA) for lympho-depletion^5,6^. Adoptive transfer of CAR-T_*shTet2-C57*_ library or control T cells transduced with truncated CAR alone into C57BL/6 mice did not induce observed toxicity (fig. S6a). Relative to control T cells, CAR-T_sh*Tet2*-C57_ library shows similar tissue distribution in mice (fig. S6b). Serum chemistry of different CAR-T cell-treated mice indicated normal pancreatic, kidney, and parathyroid functions (fig. S7a). Moreover, no significant difference of histologic changes in the main organs between different CAR-T cell treatments (fig. S7b), supporting the notion that CAR-T library constructed form naïve mouse antibody repertoires has no undesired toxicity in our study.

We next investigated in vivo anti-tumor effects of CAR-T_sh*Tet2*-C57_ library in mouse models using MC38 colorectal adenocarcinoma (Fig. 6a). MC38 tumor cells expressed high levels of PD-L1 and is a solid model to simulate the tumor microenvironment. We inoculated mice with MC38 tumors in the flank and, after tumors were palpable, treated mice with 200 mg/kg CPA and un-transduced cells, T_sh*Tet2*_ cells, or CAR-T_sh*Tet2*-C57_ library. CPA pretreatment and pretreatment with un-transduced cells or T_sh*Tet2*_ Cells resulted no inhibition of tumor growth compared with vehicle group. No reduced growth of MC38 colorectal adenocarcinoma was observed until about 30 days later, treatment with CAR-T_sh*Tet2*-C57_ library shows a significant tumour inhibition in 7 of 8 mice, and the treatment induce complete regression of tumor, which led to long-term survival of the mice without tumor recurrence, suggesting the enrichment of MC38-specific CAR-T cells exerting antitumor immunity. To evaluate if CAR-T_sh*Tet2*-C57_ library treatment confers long-term anti-cancer protection and immunological memory, complete responders of the CAR-T_sh*Tet2*-C57_ library treatment were rechallenged with MC38 cells 14 weeks after the T cell injection. All of the rechallenged mice shows no sign of tumor growth whereas as expected robust MC38 tumor growth were observed in the control mice (Fig. 6b). We further use B16.F10 melanoma model to access the anti-tumor effect of CAR-T_sh*Tet2*-C57_ library in our study (Fig. 6c). B16 tumor implants with aggressive tumor growth in all treatment group of mice, however, treatment with CAR-T_*shTet2-C57*_ cell library shows a tumour inhibition at about 40 days in 6 of 8 mice. To evaluate if CAR-T_sh*Tet2*-C57_ library treatment have ability to confer anti-cancer effect to a mutated or evolved tumour, the complete responders of N16 tumors were rechallenged with B16 cells (Fig. 6d) or MC38 cells (fig. S8), respectively. Interestingly, not only B16 cells, but also MC38 cells shows no sign of tumor growth in the rechallenged mice.

**Figure 6.**
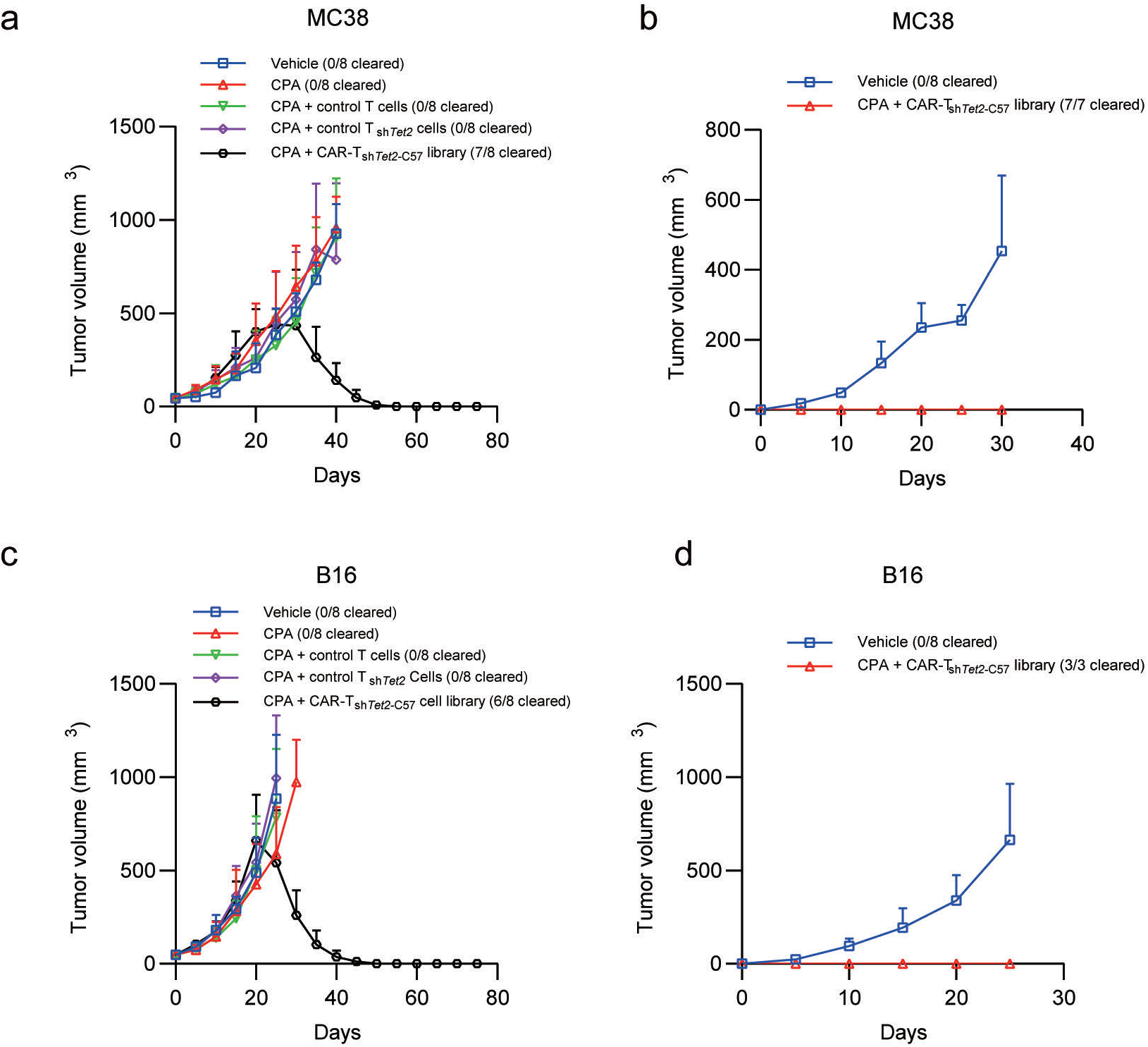
anti-tumor effect of chimeric antigen receptor T cell library in syngeneic tumor models. **a.** Tumour volumes of MC38 tumour xenografts after the indicated treatment. **b.** rechallenge of MC38 tumour xenografts with indicated treatment. **c.** Tumour volumes of B16 tumour xenografts after the indicated treatment. **d.** rechallenge of B16 tumour xenografts with indicated treatment.

### Developed of logic-gated synthetic receptor NK cell library

We next aimed to develop a large-scale CAR library platform using NK-92 cells. Similar CAR library design was used and a lentiviral vector encoding a Gal4-KRAB transcription inhibitor under the control of a NFAT-RE promoter was further constructed (Fig. 7a), and an inducible caspase-9 suicide gene was placed downstream of the Gal4-KRAB transcription inhibitor under the control of a combined UAS-SV40 promoter. Because the KRAB protein has been demonstrated to be capable of inhibiting all promoters within at least 3 kB ^16^ and because the forward construct would therefore inhibit the NFAT-RE promoter, we generated the opposite construct, in which the two expression cassettes were cloned in such a manner that both promoters were at the opposite ends and at a long distance (Fig. 7a). We postulate that, a particular antigen can activate a specific CAR, enabling that specific cell to survive and multiply under the selection pressure caused by iCAS9 inducer, therefore the specific CAR automatically be enriched. We then sought to determine the kinetics of the genetic circuit in which NFAT was coupled with KRAB activation in engineered NK-92 cells after CAR stimulation. Therefore, we used engineered NK-92 cells with either an EGFR-specific or a HER2-specific CAR (termed as NK-92 CTX or NK-92 TTZ) in our experiments. Specific cell killing effect was confirmed using standard ^51^Cr-release assays with MCF-7 and derivative cells (Fig. 7b). Then, we performed coculture experiments with the engineered NK-92 cells and different MCF-7 stimulator cells to determine whether treatment with the small molecule dimerizer AP1903 after different time would affect the apoptosis rate of the engineered T cells. Our data show that the small molecule dimerizer AP1903 had no effect on the viability of the NK-92 cells, but caused significant apoptosis/necrosis of the engineered NK cells with no stimulatory cells (Fig. 7c). However, after 24 hours of stimulation with specific antigens, respectively, the engineered T cells were notably affected by the reduced pro-apoptotic effect of AP1903, and both the EGFR-specific and HER2-specific CAR T cells were resistant to AP1903 after antigen stimulation for 72 hours (Fig. 1d). To further explore the growth dynamic of logic gated NK-92 cells, a similar small library of 10 CAR constructs comprising antigen-specific scFv were constructed, termed as the NK-92_CTX-TTZ_ library. Our data shows that repeat stimulations with MCF7 EGFR cells or MCF-7 HER2 cells resulted in no significant change of CAR composition CAR-T_CTX-TTZ_ library co-cultured with either MCF-7 cells or antgen positive MCF-7 derivatives, while enrichment of NK-92 CTX and NK-92 TTZ cells were observed post AP1903 treatment in the presence of specific MCF-7 derivatives (Fig. 7e). These data suggested the gene circuit was effective in the cell based CAR screen method, and adding of AP1903 is essential for the enrichment of CAR NK-92 cells. The antitumor effect of the CAR NK-92_CTX-TTZ_ library were further tested in immunodeficient MCF-7, MCF-EGFR and MCF-HER2 xenograft NSG mice. Mice were treated with NK-92_CTX-TTZ_ library (1×10^6^ cells for each CAR construct containing NK-92 cells). One week later, mice were treated with three doses of AP1903 intraperitoneally (i.p.) every two days as indicated (fig. S9). Notably, treatment of CAR NK-92_NSG_ library result an inhibition of tumour growth in approximately two weeks after cell injection and a complete response to the treatment was observed MCF-7 EGFR tumours or the MCF-7 HER2 tumours, suggested the expansion of the target-specific CAR-NK cells in these models.

**Figure 7.**
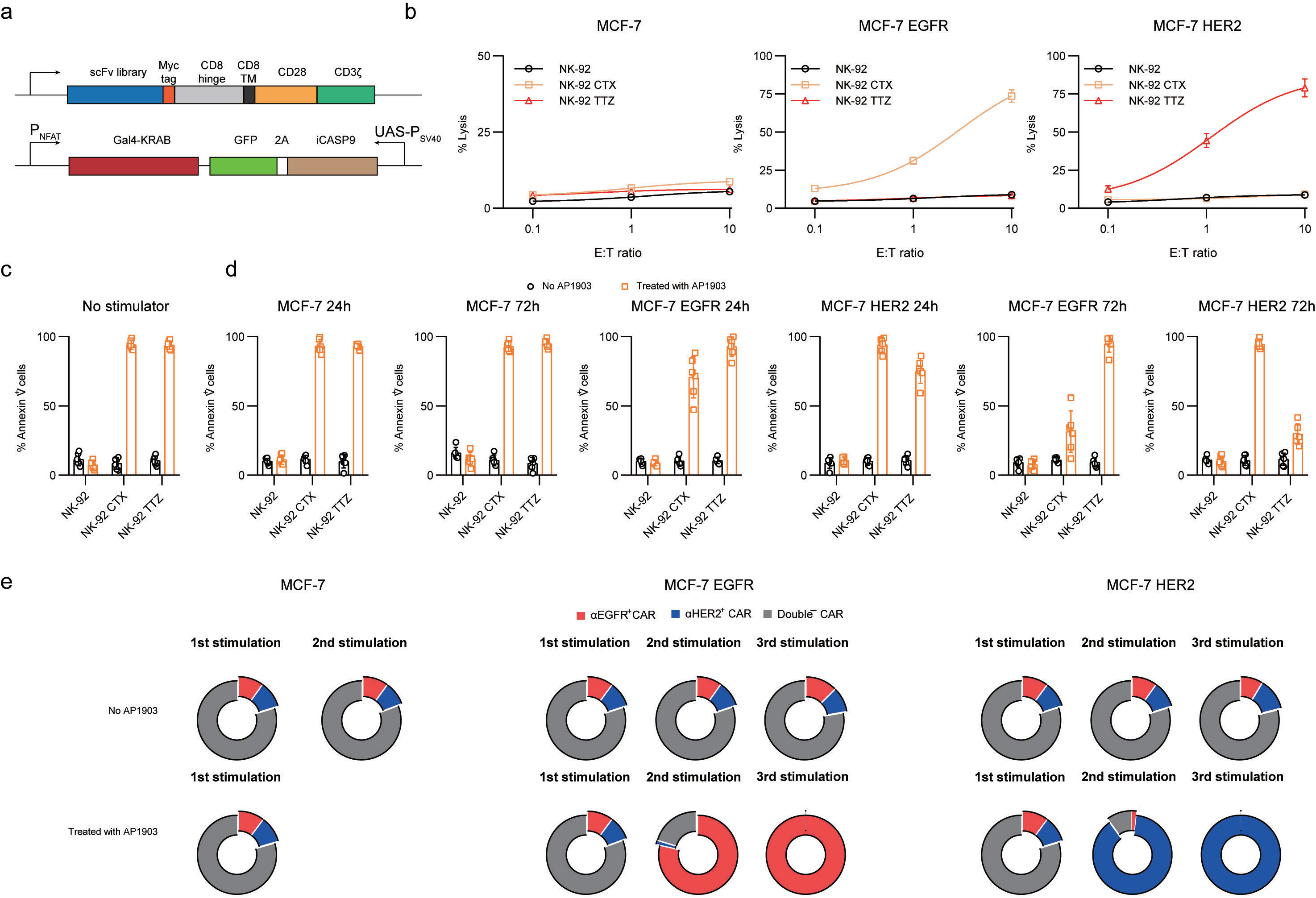
Design of a logic-gated CAR NK cell library. **a.** Map of lentiviral constructs encoding the CAR library. **b.** Killing activity of the CAR NK-92 cells in response to tumour cells. The cytotoxic activity of the CAR NK-92 and control cells against MCF-7 cells was assessed by a ^51^Cr-release assay at the indicated effector-to-target (E:T) ratios. **c and d.** cells were first stimulated with different MCF-7 cells or not for 24 or 72 hours. The addition of 10 nM AP1903 to the cultures induced apoptosis/necrosis of the CAR-NK cells, as assessed by annexin-V staining in 4 independent experiments. The dimerizer did not induce **e.** Growth dynamics of CAR-NK library in the different co-culture methods

To explore the boarder anti-tumour potential of CAR library based on NK-92 cells, a CAR NK-92 cell library containing 1×10^6^ individual CAR clones, termed as CAR NK-92_NSG_ library. Three cancer cell line xenograft models were used to test the efficacy of the CAR NK library based immunotherapy (fig. S10a). We observed a nearly complete response and a complete response to the treatment of CAR NK-92_NSG_ library in SW480 tumours and SK-OV-3 tumours, while a partial response in SK-BR-3 tumors (fig. S11). After the experiments, all the SK-BR-3-free mice were sacrificed, and the CAR^+^ NK-92 cells were further sorted to study the treatment of a second cohort of established tumours (termed CAR NK-92_SKBR3_ cells) (Fig. S10b). The SK-BR-3 xenograft mice remained untreated until tumours of approximately 500 mm^3^ size formed, and then, they were treated with 5× 10^5^ CAR NK-92_SKBR3_ cells. Our data showed that the treatment of CAR NK-92_SKBR3_ cells significantly inhibited the growth of established tumours, and all the tumours were eliminated after 3 weeks of treatment. Moreover, in both N3 and BN16 PDX models, size of tumour xenografts of control NK-92 treated mice consistently increased. The growth rate was similar to that of the xenograft of nontreated mice. In contrast, tumour xenograft size decreased in mice treated with CAR NK-92_NSG_ library (fig. 11), and complete tumour elimination was achieved in five of eight N3 tumours and six of eight BN16 tumours after 1 months of NK cell infusion.

### Expanding the therapeutic role of the CAR Repertoire to include Non-malignant diseases

To further investigate the therapeutic role of the synthetic receptor repertoire, we tested whether the CAR NK-92_NSG_ library could prevent the formation of endometriosis in vivo. Human endometrial tissue obtained from women undergoing surgery for benign conditions was implanted in NSG female mice. The animals were distributed into vehicle, control NK-92 cells or CAR NK-92_NSG_ library was administered twice for one week.

After 45 days, the human endometrial implants were excised. For the CAR NK-92_NSG_ library-treated animals, all the parietal peritoneum in which the endometrial tissue had been implanted was fully extracted because macroscopic lesions were not identified. The results of the observations are listed in Table S2.

## Discussion

Cancer remains a leading cause of mortality and poor health worldwide^17^. Despite the enormous effort expended in the development of therapeutics, successful control and eradication of the disease remain elusive^18,19^. In addition, risk factors such as physical inactivity, obesity, and smoking have shown increased prevalence worldwide, promoting the incidence of malignancy^20,21^. Moreover, our understanding of cancer genetics and biology has also been updated: cancer progression and development are evolutionary processes driven by somatic cell mutations and subclonal selection^22^, a ground-breaking theory developed by Peter Nowell and Macfarlane Burnet. Their unique theories have explicit parallels to Darwinian natural selection, in which cancer is viewed as an asexual process that produces single mutated cells and quasi-species ^22,23^. Modern cancer genomics and biology have validated the concept that cancer development is a complex, adaptive, and Darwinian process.

In humans, hundreds of millions of different antibodies (or immunoglobulin-like proteins) are created by the immune system. Antibody display technologies using phage, ribosomes, yeast and mammalian cells, have become mainstream antibody and protein engineering platforms and constitute major technology for antibody discovery and therapeutic use.

Antibody libraries can be constructed based on different sources, such as immunized animals^24^ or naturally immunized or infected humans^25^, or naive immune systems, which can be derived from nonimmune natural or computational and synthetic sources. When the diversity is sufficiently large, the library has the capacity to generate antibodies against a large number of different antigens, including self, nonimmunogenic and toxic antigens, and for this reason, these libraries are now extensively used in industry and academia^26,27^. Usually, panning for antibodies to a specific antigen using nonimmune libraries or based on synthetic antibody libraries that are diversified in merely one or two CDRs or in only one of the two chains results in the discovery of medium-affinity antibodies, reflecting the situation of transgenic mice with restricted antibody repertoires^28,29^. Interestingly, CARs constructed with high-affinity antibody fragments recognize targets expressed at any level, including in normal cells in which they were undetectable by flow cytometry, while CAR-T cells harbouring medium-affinity CAR exhibited robust antitumour efficacy similar to that exhibited by high-affinity cells but normal cells were spared from expressing physiologic target levels^30^. These findings suggested that a synthetic cell repertoire may not need the most successful antibody libraries to display (natural or synthetic) diversity in multiple CDRs and routinely yield single-digit nanomolar and sometimes sub-nanomolar affinity antibodies—the latter having affinities equal to the affinities of antibodies regularly isolated from immunized mice or from recombinant immune libraries. In general, the size of the library is proportional to the antibody affinities: up to 10 nM for libraries with 10^7^ to 10^8^ clones and up to 0.1 nM for the best libraries with over 10^10^ members. Using such libraries, thousands of antibodies that bind distinct epitopes on the same target antigen can be retrieved with high-throughput screening ^31,32^. Therefore, a synthetic cell repertoire with 10^6^ clones should be a very successful cell repertoire for therapeutic purposes.

In this report, we postulate that the mating of the mammalian-cell antibody display library builds a novel ‘molecular evolution machine’ that can function in vivo. With chimeric antigen receptors and immune cells, the new platform can be directly selected and screened for therapeutic effects, not merely binding affinity. Moreover, the whole human antibody repertoire can also be preserved when one is healthy and use it to develop a synthetic immune cell library for therapeutic use when need. This definition of the “Synthetic immunity” may represent the promised ‘land’ that Paul Ehrlich wrote about over 100 years ago^33^: “the land which […] will yield rich treasures for biology and therapeutics.

## Methods

### Cell lines

MCF-7, SW480, SK-BR-3, SK-OV-3, and 293T cell lines were purchased from the American Type Culture Collection (ATCC, Manassas, VA). The identities of the cell lines were verified by STR analysis, and the cell lines were confirmed to be mycoplasma free. The cells were maintained in DMEM with 10% foetal bovine serum. Cell culture media and supplements were obtained from Life Technologies, Inc. Human NK-92 cells (ATCC) were propagated in X-VIVO 10 medium (Lonza) supplemented with 8% heat-inactivated human plasma (German Red Cross Blood Donation Service Baden-Württemberg–Hessen, Frankfurt, Germany) and 100 IU/ml IL-2 (Proleukin; Novartis Pharma, Nürnberg, Germany). Residues 1–619 of the extracellular domain of EGFR (EGFR-ECD) and residues 1–646 of HER2-ECD were prepared using the pcDNA3.4 expression vector (Invitrogen) and FreeStyle 293 expression system (Invitrogen).

### Vector construction

The sequence encoding the scFv antibodies generated from cetuximab, trastuzumab, CH65, 9.8B, 2F5, F10, 7D11, 8D6, omalizumab, TE33, R10 and HC33.8 was chemically synthesized. The sequence information can be found in the Protein Data Bank. The 4D5-5 scFv derived from trastuzumab was chosen for this study, as we reported previously^13^. As shown in Fig. 1A, the CAR design contained the human CD8α signal peptide followed by the scFv linked in-frame to the hinge domain of the CD8α molecule, transmembrane region of the human CD8 molecule, and the intracellular signalling domains of the CD137 and CD3ζ molecules. . Unpaired cysteine 164 within the CD8a hinge region was replaced with a serine to increase CAR expression, as reported previously^34^. The constructs were further cloned into the pHR vector backbone under the control of a PGK promoter. To generate CAR response elements, a CAR inducible promoter containing six NFAT-REs in a minimal IL-2 promoter^35^ was placed upstream of the transcription factor Gal4-KRAB. Moreover, the inducible caspase-9 suicide gene that induces apoptosis upon specific binding with the small molecule dimerizer CID AP1903 was placed downstream of the expression under the control of the combined SV40/UAS promoter. Because the KRAB protein has been demonstrated to be capable of inhibiting all promoters within at least 3 kB ^36^ and because the forward construct would therefore inhibit other promoters, we generated the opposite construct, in which the two expression cassettes were cloned in a manner such that both promoters were at the opposite ends at a distance greater than 4 kb. The 2A peptide sequence was intercalated between iCASP9 and the GFP tag.

synNotch receptors and response elements were obtained from Addgene (Addgene plasmids #79123 and 79125). The ScFv library with the N-terminal CD8α signal peptide was fused to the synNotch-Gal4VP64 receptor backbone (Addgene plasmid #79125) in place of the CD19-specific scFv. To generate the response elements, the mCherry gene segment in Addgene plasmid #79123 was replaced with the described anti-EpCAM CAR transgene. To express the TetR-KRAB fusion protein concomitantly with CAR, the 2A peptide sequence was intercalated between the two genes. The ICASP9 and GFP fusion construct was placed downstream of the expression under the control of the Tet-cytomegalovirus (CMV) promoter^36^.

### In vitro T cell transduction and cultures

Negative selection using RosetteSep kits (Stem Cell Technologies) was adopted to isolate primary human T cells from healthy volunteer donors following leukapheresis. All specimens were collected under an approved protocol by the Second Military Medical University Review Board, and written informed consent was obtained from each donor. Bulk primary human T cells were activated with paramagnetic beads coated with anti-CD3 and anti-CD28 monoclonal antibodies as previously described^13^ and transduced with lentiviral vectors encoding the indicated CAR, genetic circuits or shRNA hairpin sequences targeting *TET2* or a scrambled control coexpressing BFP (GeneChem). Following 9-10 days in culture, the T cells were FACS sorted to >95% purity with the following markers: untransduced T cells: myc^−^; CAR-T cells: myc^+^; logic-gated CAR-T cells: myc^+^GFP^+^; and logic-gated CAR-T_*shTET2*_ cell: myc^+^GFP^+^BFP^+^. Sorted T cells were subsequently expanded for 3 days in medium (RPMI 1640 with 10% human serum, 2 mM L-glutamine, 25 mM HEPES, penicillin/streptomycin (100 U/ml), and 50 mM b-mercaptoethanol (Sigma)) with 50 U/ml recombinant human IL-2 (Prometheus) with 50 U/ml human IL-2 prior to in vitro assays or adoptive transfer. Knockdown efficiency in the T cells following shRNA transduction was determined by real-time quantitative PCR with TaqMan gene expression assays (Applied Biosystems) for *TET2* (assay Hs00325999_m1) and *GAPDH* (assay (Hs03929097_g1), which served as a loading and normalization control.

### Antibody library

The protocol for antibody library construction was described in our previous report^37^. Briefly, peripheral blood mononuclear cells were isolated from blood. Total RNA was extracted, and nested PCR was used to clone genes with single domain antibodies consisting of the heavy chain and light chain variable domains. The phagemid vector pCANTAB5E (GE Healthcare) carried the final PCR products and was introduced into electrocompetent *Escherichia coli* TG1 cells that were freshly prepared. The cells were selected on lysogeny broth agar plates supplemented with ampicillin and glucose cultured overnight at 37°C. After being scraped from the plates, the colonies were stored at −80°C in lysogeny broth supplemented with 20% glycerol.

### Cytotoxicity assays

The cytotoxicity of the CAR-expressing T cells or exosomes was tested in a standard 4-h ^51^Cr-release assay^13^. Target cells were labelled with 51Cr for 1 h at 37°C. Radioactive ^51^Cr (50 μCi) was used to label 1 × 10^6^ target cells. One hundred microliters of labelled target cells (n = 5000) was plated in each well of a 96-well plate. Effector cells were added at a volume of 100 μl at different E:T ratios. Exosomes were added at different concentrations. The CAR-T cells or exosomes and targets were incubated together for 4 h at 37 °C. The supernatant (30 μl) from each well was collected and transferred to the filter of a LumaPlate. The filter was allowed to dry overnight. The radioactivity released into the culture medium was measured using a β-emission-reading liquid scintillation counter. The percentage of specific lysis was calculated as follows: (sample counts – spontaneous counts)/(maximum counts – spontaneous counts) × 100.

### Cell growth assays

The cells were washed and labelled with alamarBlue (Invitrogen), and fluorescence was read using a 96-well fluorometer with excitation at 530 nm and emission at 590 nm. The results are expressed in relative fluorescence units (RFUs) and compared with the data obtained on the first day.

### Activation of the suicide gene in vitro and validation in vivo

The small molecule dimerizer AP1903 (10 nM, MCE) was added to the indicated cell cultures. The extent to which the transduced cells were eliminated was evaluated by Annexin-V staining. The efficacy of the suicide gene was determined in vivo by treating the tumour-bearing mice that had received synthetic cell treatment with the indicated doses of AP1903 (50 μg each) intraperitoneally (i.p.)^38^.

### In vivo study

In vivo experiments were approved by the Institutional Animal Care and Use Committee (IACUC) of Second Military Medical University, and the mice were housed in a specific pathogen-free barrier facility. The cancer cells were inoculated into NSG mice. (Shanghai Model Organisms Center, Inc.) When the tumour volume reached an average of approximately 50 mm^3^, the mice were randomly divided into groups of 8 mice.

The mice were injected i.p. with 200 mg/kg cyclophosphamide to further deplete the host lymphocyte compartments. The tumours were measured with digital callipers, and the tumour volumes were calculated by the following formula: volume = length × (width)^2^/2. For cell-based therapy, the mice were injected intravenously with the indicated dose or 1× 10^7^ control cells or cell libraries twice per week for one week, and one week later, three doses of AP1903 were administered to the mice treated with cell library every two days to eliminate the non-activated synthetic cells in all experiments unless otherwise specified. Mice were injected i.p. with 2000 units of IL-2 twice a week following infusion of the T cell-based therapy. The mice were sacrificed when the volume of the tumours reached 1300 mm^3^.

For the experimental model of endometriosis, the menstrual endometrium was obtained by aspiration using Cornier cannula for endometrial biopsy during the proliferative phase (from 5 to 10 days) from 6 women undergoing surgery for benign conditions (3 myomas and 3 simple cysts). The endometrium samples were preserved in physiological saline and implanted immediately to prevent cell death. The patients were of reproductive age (23-45 years old) and had not received hormone treatment within 8 weeks prior to sample collection. The medical history of endometriosis or adenomyosis was not available; however, an extensive preoperative evaluation was performed for all the patients, and it included clinical exploration, MRI, and transvaginal sonography to exclude both endometriosis and adenomyosis. The absence of endometriotic lesions was confirmed during laparoscopy conducted for the determination of benign disease. All specimens were collected under an approved protocol by the Second Military Medical University Review Board, and written informed consent was obtained from each donor.

### Statistical analysis

Unless otherwise specified, Student’s t-test was used to evaluate the significance of differences between 2 groups, and ANOVA was used to evaluate differences among 3 or more groups. Differences between samples were considered significant when *P* < 0.05.

