## Supplementary figures and tables for "Synthetic Cellular Immunity based on CAR repertoire for Human Immunotherapy"

##### *External Data*

**Contents** **Page**

### Supplementary Figures

Figure S1

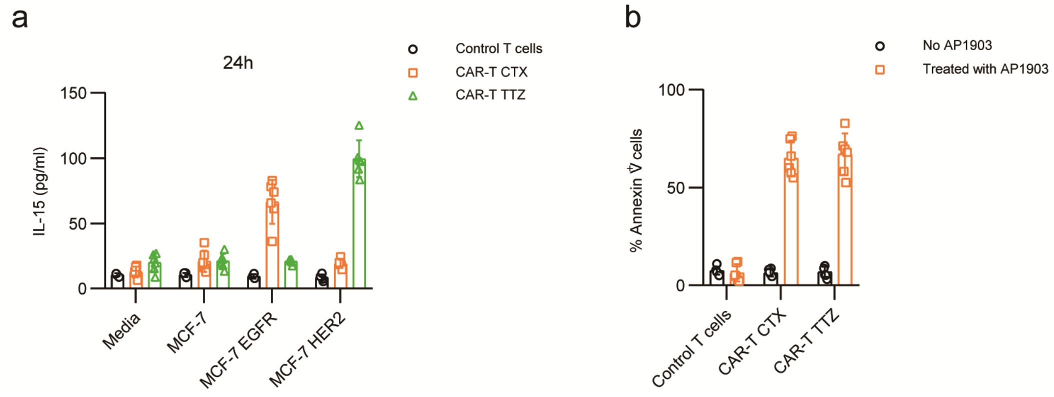

**Figure S1. Vector validation. a**. IL-15 reponse to CAR activation. **b**. The addition of 10 nM AP1903 to the cultures induced apoptosis/necrosis of the CAR-T cells

Figure S2

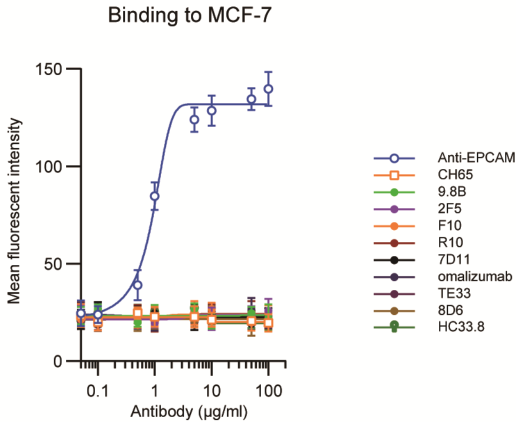

**Figure S2. Binding of scFvs to MCF-7 cells.** Different scFvs were evaluated for their ability to binding to MCF-7 cells.

Figure S3

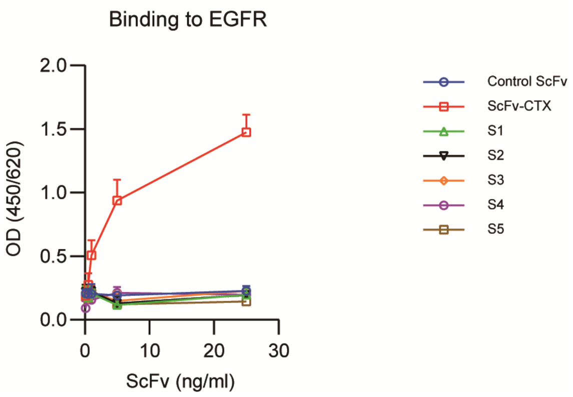

**Figure S3. Binding of scFvs to EGFR.** Different scFvs were evaluated for their ability to binding to EGFR.

Figure S4

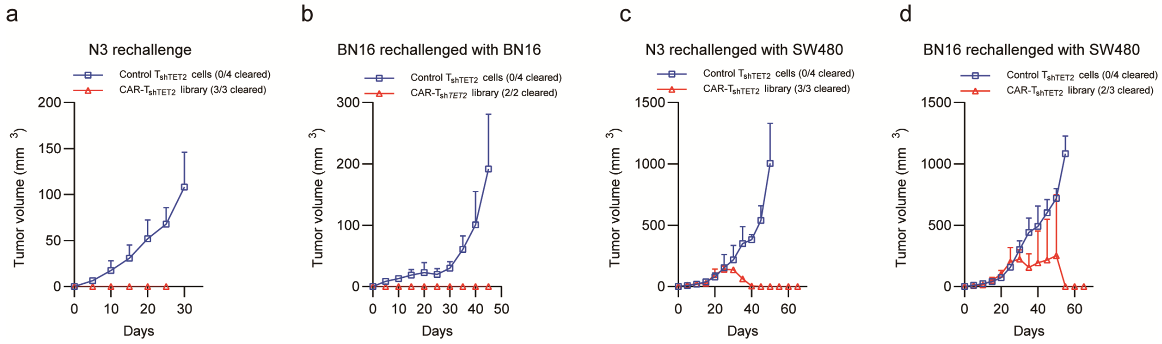

**Figure S4.Tumor model rechallenge.** Tumour volumes of different tumour xenografts after the indicated treatment.

Figure S5

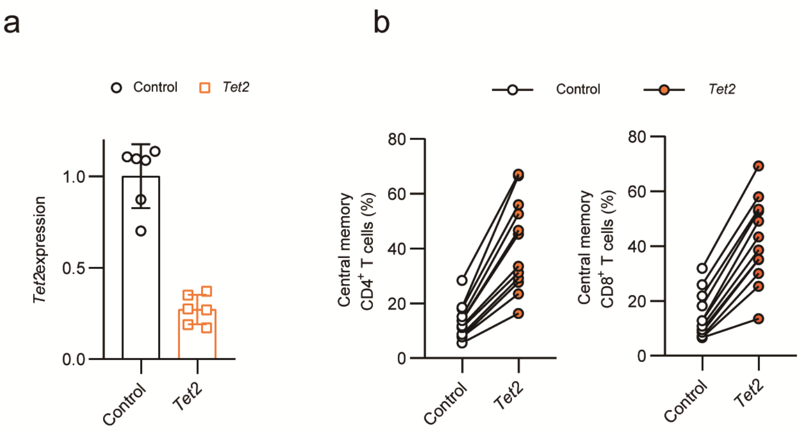

**Figure S5.murine Tet2 modified cells.**

Figure S6

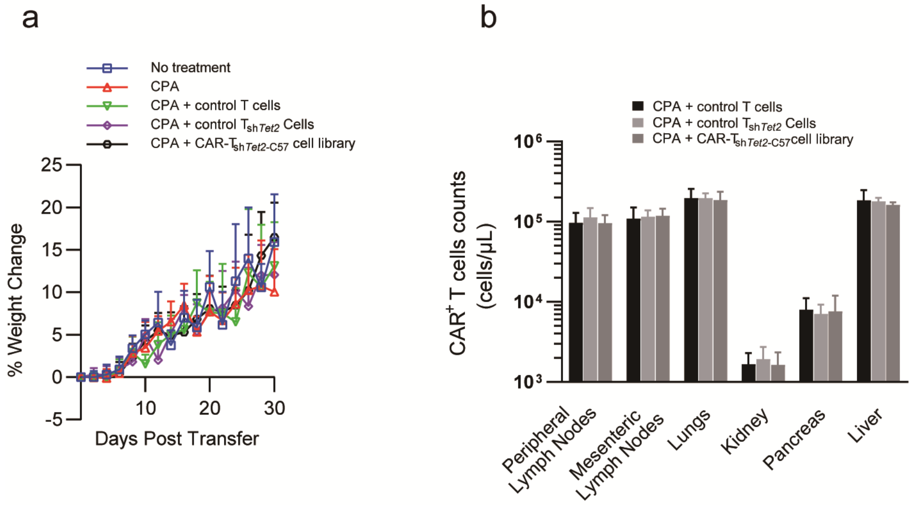

**Figure S6. CAR-T Cell library did not induce toxicity**

Figure S7

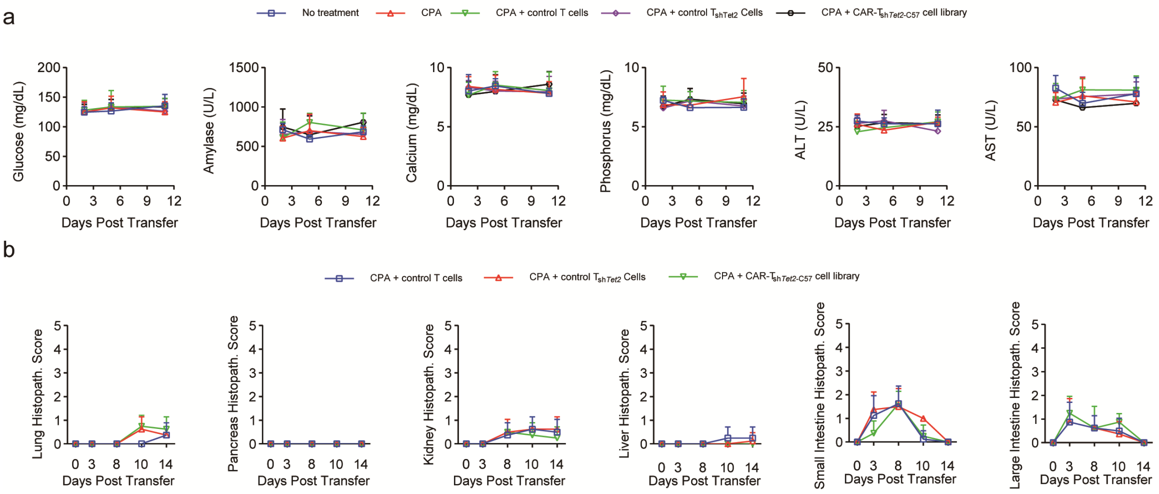

**Figure S7. Serum chemistry analysis and Histopathology scoring**

Figure S8

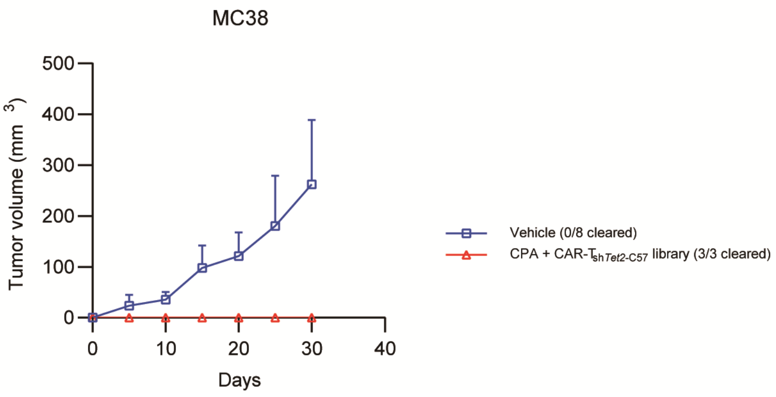

**Figure S8.Tumor model rechallenge.** Tumour volumes of different tumour xenografts after the indicated treatment.

Figure S9

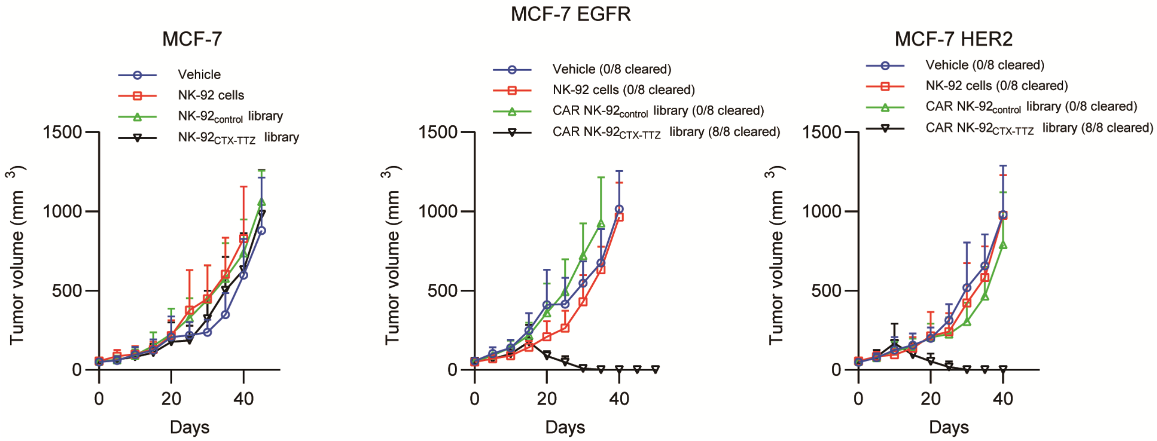

**Figure S9.Anti-tumor effect of NK92 cell library.** Tumour volumes of different tumour xenografts after the indicated treatment.

Figure S10

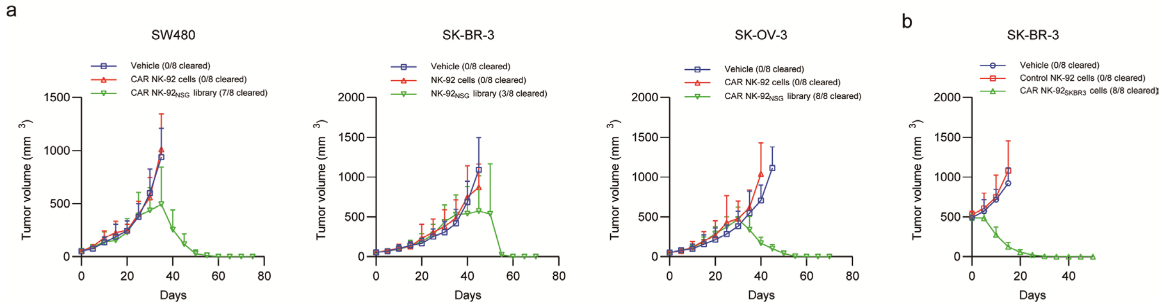

**Figure S10.Anti-tumor effect of NK92 cell library.** Tumour volumes of different tumour xenografts after the indicated treatment.

Figure S11

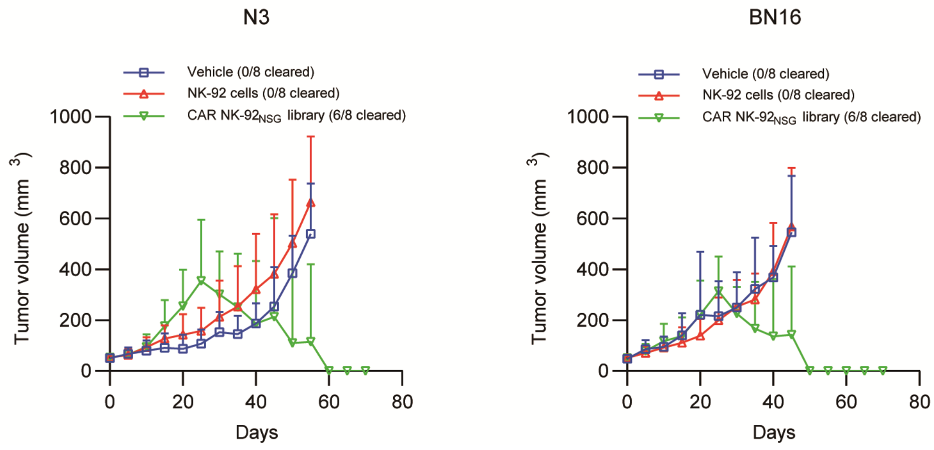

**Figure S11.Anti-tumor effect of NK92 cell library in PDX models.** Tumour volumes of different tumour xenografts after the indicated treatment.

### Supplementary Tables

#### Table S1. Control antibodies used in the study

| Antibody name | Reference (PMID) |
|---|---|
| CH65 | 21825125 |
| 9.8B | 23577234 |
| 2F5 | 19740978 |
| F10 | 19234466 |
| 7D11 | 17688903 |
| 8D6 | 30069035 |
| omalizumab | 26113483 |
| TE33 | 17712773 |
| R10 | 28074040 |
| HC33.8 | 26739044 |

**Table S2. Macroscopic Results of Endometriosis-Like Lesions According to Treatment Groups**

| Measurements | Vehicle | NK-92 cells | CAR NK-92$_{NSG}$ library |
|---|---|---|---|
| # of implants (n) | 2.46 ± 0.52 | 2.2 ± 0.77 | 0 |
| Tissue volume (mm$^3$) | 424.34 ± 161.75 | 375 ± 152.23 | 0 |
